# Consequences of population structure for sex allocation and sexual conflict

**DOI:** 10.1101/2020.04.16.042994

**Authors:** Leonor R. Rodrigues, Mario Torralba Sáez, João Alpedrinha, Sophie Lefèvre, Muriel Brengues, Sara Magalhães, Alison B. Duncan

**Affiliations:** cE3c: Centre for Ecology, Evolution, and Environmental Changes, Faculty of Sciences, University of Lisbon, Edifício C2, 38 piso, 1749-016 Lisboa, Portugal; Institut des Sciences de l’Évolution, Université de Montpellier, CNRS, IRD, EPHE, CC065, Place Eugène Bataillon, 34095 Montpellier Cedex 05, France; IRCM, INSERM, Univ. Montpellier, ICM, Montpellier, France

**Keywords:** local mate competition, hard and soft selection, experimental evolution, budding dispersal, scale of competition, *Tetranychus urticae*

## Abstract

Both sex allocation and sexual conflict can be modulated by spatial structure. However, how the interplay between the type of dispersal and the scale of competition simultaneously affects these traits in sub-divided populations is rarely considered.

We investigated sex allocation and sexual conflict evolution in meta-populations of the spider mite *Tetranychus urticae* evolving under budding (pairing females from the same patch) or random (pairing females from different patches) dispersal and either local (fixed sampling from each subpopulation) *versus* global (sampling as a function of subpopulation productivity) competition.

Females evolving under budding dispersal produced less female-biased offspring sex ratios than those from the random dispersal selection regimes, contradicting theoretical predictions. In turn, the scale of competition did not have a strong effect on sex allocation. Males evolved under budding dispersal induced less female harm than those exposed to random dispersal, but there was no reduction in female fitness following exposure to multiple mates from either selection regime.

This work highlights that population structure can impact the evolution of sex allocation and sexual conflict. We also discuss how selection on either trait may reciprocally affect the evolution of the other, for example via effects on fecundity.

## Introduction

Many organisms exist in structured populations, sub-divided into patches, that are linked and shaped by demographic factors such as dispersal. The frequency and type of dispersal can determine whether interactions are more likely to occur among related or unrelated individuals (Hamilton 1964; Bulmer 1986; Queller 1992; Courteau and Lessard 2000; Rousset 2004; West 2009). For instance, if dispersal is limited, such that only some individuals disperse, the probability of interactions among genetically related individuals in a patch increases compared to populations in which all individuals disperse (Hamilton 1964; Taylor 1992; Wilson et al. 1992; Taylor and Crespi 1994). However, if individuals disperse in groups from the same patch (i.e., if there is budding dispersal), interactions among genetically related individuals can be maintained, even if dispersal rates are high (Aviles 1993; Gardner and West 2006; Lehmann et al. 2006; Gardner et al. 2009; Lehmann and Rousset 2010). Dispersal frequency and timing also influence the scale of competition. For example, high dispersal, and dispersal occurring prior to the competitive interaction, favour global competition, in which individuals compete with equal probability with others in the population (Taylor 1992; West et al. 2002a; Griffin et al. 2004). In contrast, limited dispersal, and/or dispersal occurring after the competitive interaction, favour local competition (i.e., competition within the natal patch) (Taylor 1992; Wilson et al. 1992; Frank 1998; West et al. 2002a; Griffin et al. 2004). Therefore, the type, frequency and timing of dispersal can have a significant impact on relatedness within a patch and the scale at which competitive interactions occur.

In turn, both relatedness and the scale of competition can affect sex allocation - the differential investment into male and female offspring. Indeed, in subdivided populations, sex allocation theory predicts an offspring sex-bias towards the sex for which local competition between kin is less intense (Hamilton 1967; Charnov 1982; West 2009). For example, more female-biased offspring sex ratios are predicted when males compete locally on their natal patch for mates, and mated females disperse and compete globally for new patches (Hamilton 1967; Taylor 1981; Herre 1985). If there is budding dispersal, relatedness among the offspring of foundresses increases, exacerbating local competition between related males for mates, thus selecting for even more female-biased sex ratios (Gardner et al. 2009). However, if the proportion of individuals dispersing is limited, and females compete locally for resources, competition becomes intense for both sexes and selection favours a more balanced offspring sex ratio (Table S1; Frank 1985; Herre 1985; Taylor and Crespi 1994; Courteau and Lessard 2000). A few empirical studies to date have investigated the consequences of budding dispersal (Kummerli et al. 2009), or disentangled the relative effects of the scale of competition and relatedness (Griffin et al. 2004) on the evolution of kin-selected behaviours, but none have disentangled the effect of these two factors on sex allocation.

Population structure is also predicted to impact the evolution of sexual conflict, i.e., asymmetric reproductive interests between mating partners (Bourke 2009; Pizzari et al. 2015). Theoretical work predicts that global competition selects for reduced harming behaviour of males, when interactions occur among kin, as harm reduces patch productivity (Rankin 2011; Pizzari and Gardner 2012; Pizzari et al. 2015). A number of empirical studies are compatible with this prediction (Carazo et al. 2014; Hollis et al. 2015; Lukasiewicz et al. 2017, but see Chippindale et al. 2015). For instance, in the fruit fly *Drosophila melanogaster*, females repeatedly exposed to related, as opposed to unrelated, males presented a higher lifetime reproductive success (Carazo et al. 2014). The type of dispersal may also be crucial for the evolution of sexual conflict. As described above, random dispersal reduces relatedness among competitors, which might increase the intensity of sexual conflict (Rankin 2011; Faria et al. 2015). In contrast, sexual conflict may be reduced by budding dispersal, which maintains interactions among kin.

Curiously, despite the fact that population structure is predicted to affect sex allocation and sexual conflict (Bourke 2009), no study to date has disentangled how the type of dispersal and the scale of competition impacts the evolution of both within the same set-up. This is at odds with the fact that evolution under different population structures may simultaneously impact sex allocation and sexual conflict in a non-independent manner, highlighting the need to integrate studies on these traits (Chapman 2009; Scharer and Janicke 2009). For instance, changes in sex allocation may result in the production of more or fewer individuals of each sex, which impacts sexual conflict. At the same time, sexual conflict may impact the number of offspring produced (Carazo et al. 2014; Lukasiewicz et al. 2017), which may in turn influence sex allocation (Stubblefield and Seger 1990). This is supported by studies showing that multiple mating can impede optimal sex allocation in the parasitoid wasp *Nasonia vitripennis* (Boulton and Shuker 2015; Boulton et al. 2019).

Here, we uncover the effects of the type of dispersal and the scale of competition, on the evolution of sex allocation and sexual conflict in the spider mite *Tetranychus urticae*. Previous work has shown sex allocation and sexual conflict evolution in *T. urticae* when males compete locally for mates and females compete globally for patches (Macke et al. 2011; Macke et al. 2014), and that multiple mating can be costly for females (Macke et al. 2012; Rodrigues et al. 2020). In a fully crossed design, using experimental evolution, we placed replicate populations of *T. urticae* in 4 selection regimes with either local or global competition, and random *versus* budding dispersal. This design enabled us to follow evolution of both sex ratio and sexual conflict under different population structures.

## Material and Methods

### Biological model

The two-spotted spider mite, *T. urticae* Koch (Acari: Tetranychidae), is a generalist herbivore with a host range of over 1100 plant species (Helle and Sabelis 1985; Migeon and Dorkeld 2019). *T. urticae* has an arrhenotokous haplodiploid life cycle (∼14 days egg – adult at 20-25°C): sons develop from unfertilised, haploid eggs and daughters from fertilised, diploid eggs. We report tertiary sex ratios (adult males divided by the total number of adult offspring) as males and females can only be distinguished as adults using microscopy: males are smaller than females and possess a pointed abdomen.

### Population origins

In 2013, 10 different *T. urticae* populations were collected and separately maintained on bean plants at 25 ± 2°C, with a 16h light: 8h dark cycle at the University of Lisbon. These populations comprised seven populations from Portugal (Lou, DC, AMP, DF, CH, COL and RF), two from Spain (Albe and Alro) and one from France (FR) (Zélé et al. 2018). All populations were treated with antibiotics to ensure that they were free of bacterial endosymbionts, known to be sex ratio distorters (Breeuwer 1997). The sex ratio of each individual population ranges from 0.22 to 0.40 (Zélé et al. 2020). In November 2015, more than 50 females from each of the 10 populations were transferred to the University of Montpellier and mixed to form a genetically diverse population to seed the experiment (hereafter called the ‘ancestral population’). This newly mixed population was maintained on 12 whole bean plants (variety: Pongo) in a plastic box (395 mm length x 335 mm width x 323 mm height) at 25°C with a 16h light: 8h dark cycle. Each week, old plants were removed and replaced with young, un-infested plants. All bean plants used to maintain mite populations and for all experiments described below were grown from seeds in an isolated, herbivore-free room at 23 ± 1°C with a photoperiod of 12h light: 12h dark at the University of Montpellier.

Fourteen days before starting the experiment, 10 independent groups of 40 females were haphazardly sampled from the ancestral population and put on a patch (10-15 bean leaves placed together) on water-saturated cotton wool to lay eggs. This allowed maternal effects to be equalised and ensured that females seeding the experiment were of the same age. Two weeks later, when mites of the following generation had reached adulthood, all 10 groups were mixed, and the newly emerged adult females were haphazardly assigned to the different selection regimes.

### Establishment and maintenance of selection regimes

The impact of different types of dispersal (budding *versus* random) and scales of competition (local *versus* global) on the evolution of sex allocation and sexual conflict in *T. urticae* was investigated using a fully crossed experimental design (Figure 1): 1) global competition, budding dispersal (‘Global Budding’, GB), 2) global competition, random dispersal (‘Global Random’, GR), 3) local competition, budding dispersal (‘Local Budding’, LB) and 4) local competition, random dispersal (‘Local Random’, LR). Each regime was replicated three times (GB-1, GB-2, GB-3, GR-1, GR-2, GR-3, LB-1, LB-2, LB-3, LR-1, LR-2 and LR-3).

**Figure 1.**
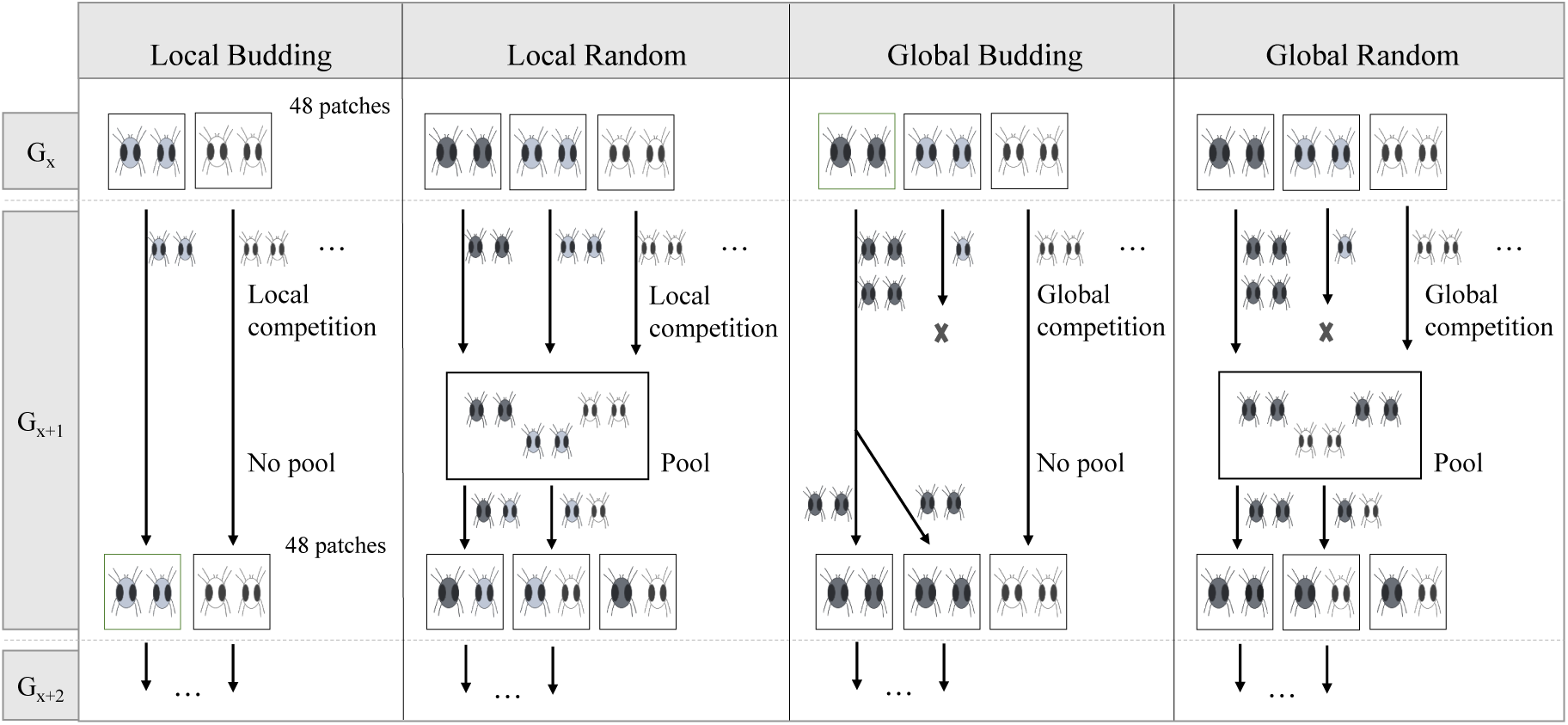
Establishment of the selection regimes. Four selection regimes were established and maintained for 33 generations, each with three experimental replicates. In ‘Local Budding’, 2 females from each of the 48 patches (squares) were transferred in pairs to a new patch for the next generation (G_x+1_). In ‘Local Random’, an equal number of females (2 – 4; the number was adjusted each generation to account for mortality) from each patch were pooled together on a large common leaf patch (‘mixing patch’, rectangle), from which females were subsequently haphazardly transferred in pairs to 48 new patches. In ‘Global Budding’, the number of adult females per patch was counted before each transfer to calculate fecundity relative to that of the other 47 patches in the replicate. Each patch contributed a number of female pairs, to the following generation, proportional to its relative fecundity. In ‘Global Random’ all 48 patches were placed on a ‘mixing patch’ onto which females could disperse for ∼4 hours, after which adult females were haphazardly transferred in pairs to 48 new patches for the next generation. Related females within a treatment are denoted as the same shade of grey.

For each replicate population, each generation comprised a total of 96 adult females, being assigned in pairs to 48 bean leaf patches (4cm^2^ each) placed on water-saturated cotton wool in a plastic box (255 mm length ×183 mm width × 77 mm height). All replicates from all regimes were maintained in the same conditions, the only difference being how populations were mixed and transferred to new patches at each generation (Figure 1).

In the budding dispersal regimes, females were always transferred with another female from the same patch to form the next generation. In contrast, in the random dispersal regimes, females from different patches were placed together on a ‘mixing patch’ (10 bean leaves placed together) before being transferred, in haphazardly chosen pairs, to a new patch. Local competition was imposed by letting an equal number of adult females per patch seed the next generation (2 – 4 females per patch in ‘Local Random’; adjusted each generation to accommodate mortality). Under global competition, relative patch productivity (the total number of daughters produced compared to that of the other patches within the replicate) determined the number of female adult offspring transferred to the next generation: in the ‘Global Random’ regime, all 48 patches were placed on a ‘mixing patch’ onto which adult females dispersed (patches with more female offspring having a higher representation on the ‘mixing patch’) before being transferred in pairs; in the ‘Global Budding’ regime, the number of adult females on each patch was counted to calculate relative fecundity (i.e. dividing the number of females per patch by the total number of females across the 48 patches), so that patches with the most offspring contributed more pairs of females to the next generation.

Due to the time taken for each transfer, transfers from one generation to the next were done over 1, 2 or 3 days. When done over more than one day, at least one replicate population from each regime was transferred on the same day. All replicates were maintained in a climate chamber at 25 ± 2°C, with a photoperiod of 16h light: 8h dark. During the selection experiment, all replicates in the ‘Local Budding’ regime were lost after generation 14, and 1 replicate in the ‘Global Budding’ regime was lost at generation 22 (GB-3). In total, 33 generations of selection were performed.

### Responses to selection

#### 1. Sex allocation during experimental evolution

The sex allocation of females was measured directly in the replicate populations of each selection regime at generations 12, 17, 20 and 31. This was done by counting the number of males and females per patch within each experimental replicate prior to the following transfer. Thus, sex ratio comprised the combined output of the two females per patch.

#### 2. Sex allocation in a common environment

In this assay, all regimes were each exposed to a common environment for 1 generation to equilibrate maternal effects before measuring the offspring sex-ratios of females that mated randomly with males from their selection regime (Figure S1). For this, at generation 31, 96 mated daughters were haphazardly chosen from the 48 patches within each selection regime and placed on a large leaf patch (∼200cm^2^) where they laid eggs together. Fourteen days later, the offspring on these patches emerged as adults and mated amongst themselves (Generation 31 + 1). Ninety-six mated female offspring from each mixing patch were then haphazardly chosen to measure their offspring sex-ratio; 48 were placed individually on 2cm^2^ patches, and another 48 placed in pairs on 4 cm^2^ patches. Females were allowed to lay eggs for 7 days on these new patches, before being killed. After 2 weeks, once offspring had emerged as adults, the number of daughters and sons on each patch was counted. This experiment was set up over three days, with one replicate per regime being treated each day.

#### 3. Sex allocation in response to patch fecundity

Measures of offspring sex ratio on patches concern the sex allocation of two females on that patch. While this is informative, it may obscure responses to selection, especially if offspring sex ratio differs between females, for instance, if a focal female’s sex allocation changes in response to her own fecundity only, or also to that of her patch mate (Stubblefield and Seger 1990). To test this hypothesis, we measured the sex allocation of single females from our selection regimes in response to the presence of eggs -that do not hatch-laid by sterile females on the same patch (Osouli et al. 2014).

This experiment was implemented after 33 generations of selection. As for the preceding experiment, individuals within each replicate population were subject to a common environment. However, in this experiment it was over two generations (generation 33 + 2; Figure S1). At the same time, females from the ancestral population were placed in a common environment for 2 generations, as done with females from the selection regimes (Figure S1) to generate sterile females. To sterilise these females, they were exposed to 100 Gy, at a dose of 2.7 Gy minute^-1^, using a Xstrahl XenX pre-clinical irradiator at the Institute of Cancer Research, Montpellier (IRCM). Preliminary studies revealed that this dose of X-ray irradiation is sufficient to sterilise females, that lay eggs that do not hatch (see Table S2).

Single adult females from the different selection regimes were placed on individual leaf patches with one sterile female and allowed to lay eggs for 5 days. Both females were then killed and the total number of eggs per patch (laid by the sterile and the fertile female, coming from the ancestral population and a selection regime, respectively) was counted. Nine days later the adult offspring of the fertile female were counted, and the offspring sex ratio measured. A total of 48 leaf patches (4 cm^2^) were set up per replicate population.

#### 4. Sexual conflict

The impact of mating with males evolved under the ‘Global Budding’ and ‘Global Random’ selection regimes on the fecundity of females from the ancestral population was compared in a separate assay. Females were collected from the different selection regimes at generation 33 and spent two further generations in a common environment before the experiment (G33 + 2, as above; Figure S1). The females from the ancestral population experienced one generation in a common environment, being placed in 2 boxes, each containing 100 females on a large ‘mixing patch’. Thirteen days later, 240 quiescent, virgin females (i.e., daughters) were isolated on 4 cm^2^ individual leaf patches later used to measure the degree of sexual conflict.

To obtain males from each selection regime, on days 10 and 11 of the second generation in the common environment (G33 + 2), 30 quiescent, juvenile females were isolated from each replicate population and each placed on a 4cm^2^ leaf patch. These virgin females emerged as adults and laid eggs for six days. Because spider mites are haplodiploid, only male progeny emerged from these eggs. Due to female mortality or failure to lay eggs, the total number of patches containing virgin males from each line varied from 17 to 28 (GB-1 = 28, GB-2 = 17, GR-1 = 21, GR-2 = 21 and GR-3 = 21). On day 1 of the experiment, males from the different patches within each replicate population were mixed on a large leaf patch so they could be haphazardly distributed across treatments (see below).

The 240 quiescent, virgin females (i.e. daughters) were taken from the ancestral population and were kept isolated for 2 days on their individual patches. Subsequently, the eggs laid by these females were removed and patches were assigned to males from either the ‘Global Random’ or ‘Global Budding’ selection regime, and to a ‘single’ or ‘double’ mate treatment (N=30 per treatment). In all treatments, males from the selection regimes were placed with the virgin females for 5 hours on day 1 of the experiment. Twenty-four hours later (day 2), in patches assigned to the double mating treatment, a second male was placed on the patch and left for 5 hours. In both treatments, females were left to lay eggs and on day 6 of the experiment, female mortality was checked and females alive were removed from the patches. The total number of eggs per patch was counted and, 8 days later, offspring sex-ratio was measured.

### Statistical analysis

All analyses were carried out using the R statistical package (v. 3.0.3) and JMP13. We used Generalised Linear Mixed Models (GLMMs, package glmmTMB; Brooks et al. 2017) with a beta-binomial error structure and logit link function, and quasi-poisson or negative binomial error structures and log link function, to analyse the effect of selection regime on sex ratio and mean offspring production, respectively. Maximal models were simplified by sequentially eliminating non-significant terms (p < 0.05) from the highest-to the simplest-order interaction, with the highest p-value to establish a minimal model (Crawley 2007). The significance of the explanatory variables in the minimal models was established using chi-squared tests (Bolker et al. 2009). *A posteriori* contrasts with Bonferroni corrections were done to interpret the effect of selection regime when significant (glht, multcomp package; Hothorn et al. 2008). Details of all models are given in Table S3.

#### 1. Sex allocation during experimental evolution

To analyse the impact of the selection regime on offspring sex ratio, generation (12, 17, 20 and 31), selection regime (GB, GR and LB) and their interaction were included in the model as fixed factors. Generation was analysed as a covariate and was log transformed to improve the fit of the model. Experimental replicate (GB-1, GB-2, GR-1, GR-2, GR-3, LR-2 and LR-3,) was included as a random factor nested within selection regime, and the day measurements were taken as a random factor nested within generation.

#### 2. Sex allocation in a common environment

To investigate the effect of selection regime on offspring sex ratio in a common environment, we used a model with selection regime (GB, GR and LB), the number of females per patch (1 or 2) and their interaction as fixed factors, and replicate population (GR-3, GB-1, GB-2, GR-1,GR-2, LR-2 and LR-3), nested within selection regime as a random factor. This analysis excluded replicate LR-1 due to fewer than 8 patches with more than 3 offspring. For this variable, the best fit model included a parameter to account for zero inflation (ziformula∼1; package glmmTMB; Brooks et al. 2017).

#### 3. Sex allocation in response to patch fecundity

In a second analysis, using data from the ‘*Sex allocation in response to patch fecundity’* experiment, we investigated whether the sex allocation of the focal female depended on her relative fecundity (‘relative patch fecundity’: the number of eggs laid by the focal female divided by the total number of eggs laid on the patch) and on the total number of eggs present in the patch (‘total patch fecundity’). In this analysis, the selection regime of the focal female (GB, GR and LB) and its interaction with relative (or total) patch fecundity were included in models as fixed factors, and experimental replicate (GB-1, GB-2, GR-2, GR-3, LR-2 and LR-3) nested within selection regime as a random factor. These analyses only included females alive on day 4 of the experiment and excluded replicates GR-1 and LR-1, due to fewer than 10 patches with more than 3 offspring.

We used data from this experiment to compare observed offspring sex ratios with predicted values from theoretical models (see Supplementary Materials Table S1 for details) using two tailed t-tests in JMP13. Observed offspring sex ratios were mean values for fertile females from each selection regime.

#### 4. Sexual conflict

To test whether selection regime affected the intensity of sexual conflict and male-male competition, female fecundity and offspring sex-ratio were analysed including the number of mates (one or two mates), the selection regime of the male (‘Global Budding’ *versus* ‘Global Random’) and their interaction as discrete, fixed variables in the model. Replicate population (GB-1, GB-2, GR-1, GR-2 and GR-3) and box (the container in which several individual replicates were maintained) were included nested within dispersal type as random factors. In the analysis of female fecundity, all individual replicates in which females died before day six were excluded.

## Results

### Sex allocation during experimental evolution and in a common environment

There was a consistent significant effect of selection regime on sex allocation during the selection experiment and after a generation in a common environment (during selection experiment: X^2^_2_ = 14.046, p < 0.001; common environment: X^2^_2_ = 11.845, p = 0.002; Figures 2a and 2b, Table S4). Indeed, females from the ‘Global Budding’ regime produced less female-biased offspring sex ratios than females from the ‘Global Random’ regime (during selection experiment: Z = -3.741, p < 0.001; common environment: Z=-3.384, p = 0.002; Table S5). There was also a trend for females from the ‘Global Budding’ regime to produce a less female-biased offspring sex ratio than females from the ‘Local Random’ regime during the selection experiment (Z=-2.289, p = 0.066), but not after a generation in a common environment (Z=-1.53, p = 0.331 Figures 2a and 2b; Table S5). There was no difference in sex allocation between females from the ‘Global Random’ and ‘Local Random’ regimes (during selection experiment: Z=1.554, p = 0.361; common environment: Z=-1.597, p = 0.3776; Figures 2a and 2b; Table S5). The number of females on a patch had no effect on offspring sex ratio (selection regime x number of females per patch: X^2^_2_ = 4.114, p = 0.128; number of females per patch: X^2^_1_ = 0.94, p = 0.331; Table S4).

**Figure 2.**
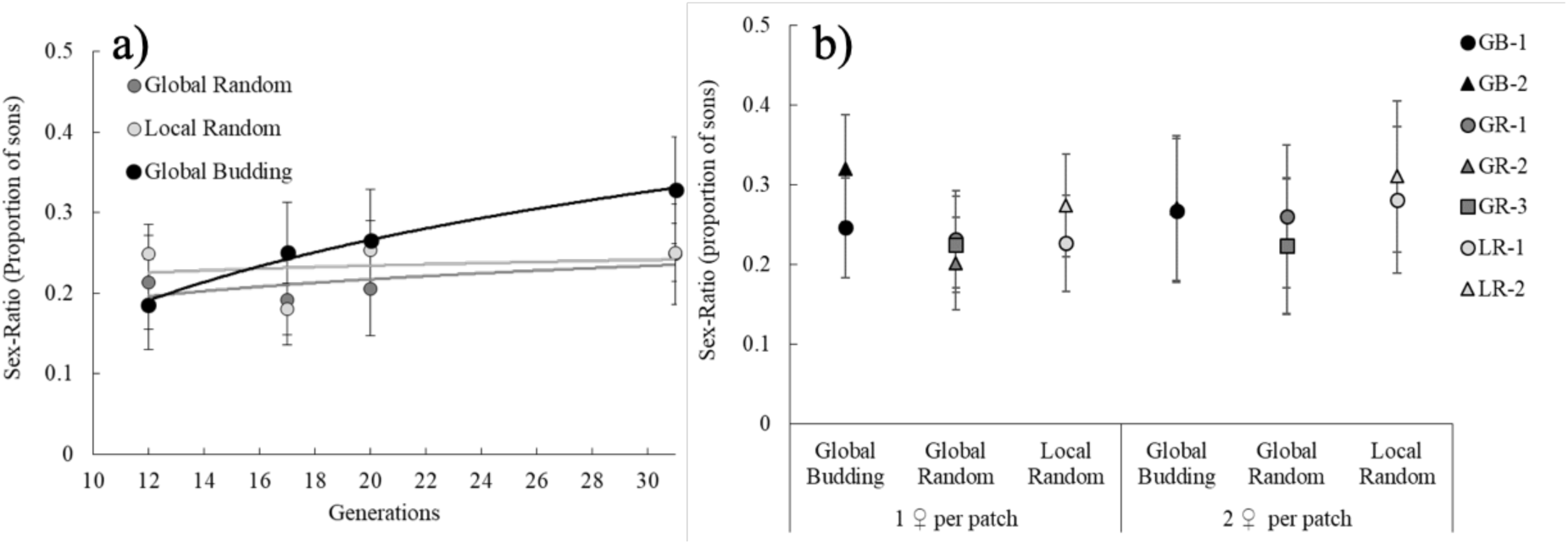
Mean offspring sex ratio (± standard error) of females from the ‘Global Random’ (GR, dark grey), ‘Global Budding’ (GB, black) and ‘Local Random’ (LR, light grey) selection regimes across generations. The proportion of male offspring was measured a) during experimental evolution at generations 12, 17, 20 and 31 (means shown for each selection regime) and b) at generation 31 + 1 after one generation in a common environment, in patches with one or two females (means shown for each experimental replicate (different symbols) in each selection regime).

### Comparing observed offspring sex-ratios with theoretical predictions

We compared mean offspring sex ratios obtained in the different selection regimes with theoretical predictions (Taylor and Bulmer 1980; Herre 1985; Gardner et al. 2009) (See Table S1 for predicted offspring sex ratios).

Females from the ‘Global Random’ selection regime produced an offspring sex ratio of 0.19 (± 0.19 SE), that does not differ from the predictions of Taylor and Bulmer (1980) and of Gardner et al (2009) (t = 0.932, df = 69, p = 0.3544). In contrast, the evolved offspring sex ratios in the ‘Global Budding’ and ‘Local Random’ selection regimes differed from theoretical predictions. Specifically, females from the ‘Global Budding’ selection regime produced a less female-biased offspring sex ratio (mean 0.30 ± 0.03 SE; t = 9.54, df = 55, p < 0.0001), and females from the ‘Local Random’ regime a more female-biased offspring sex ratio (mean 0.24 ± 0.02 SE; t = 7.99, df = 74, p < 0.0001), than predicted by theory.

### Sex allocation in response to patch fecundity

Offspring sex ratios changed according to the selection regime of the focal female and her relative patch fecundity (selection regime: X^2^_2_ = 10.90, p = 0.004; relative patch fecundity: X^2^_1_ = 6.87, p = 0.009; Figure 3, Table S4). As before, females from the ‘Global Budding’ regime produced less female-biased offspring sex ratio than females from the ‘Global Random’ regime (Z=-3.298, p = 0.003; Figure 3, Table S5). The offspring sex ratio of females from the ‘Local Random’ treatment did not differ from that of the other two selection regimes (Table S5). Across all treatments, females with higher relative patch fecundity produced more female-biased offspring sex-ratios (selection regime x relative patch fecundity: X^2^_2_ = 2.55, p = 0.28; Figure 3). These results did not change when using total patch fecundity (sum number of eggs laid by the fertile and sterile female on each patch, Figure S2, Tables S4 and S5;).

**Figure 3.**
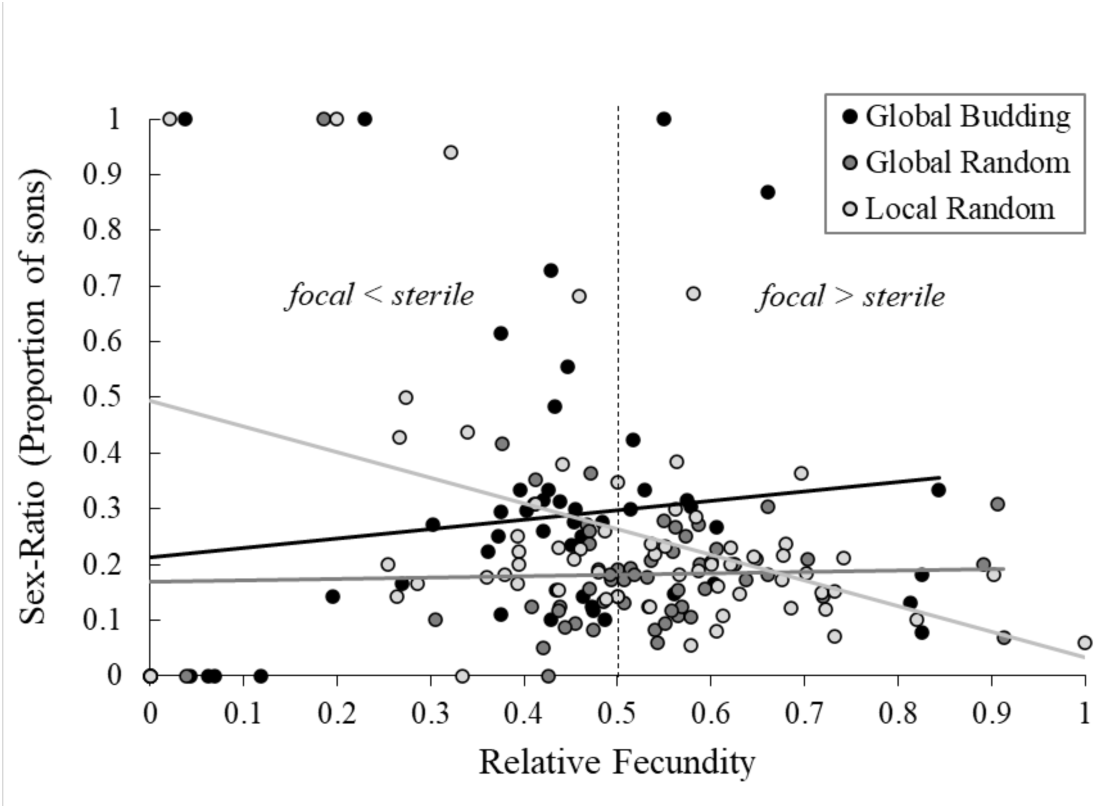
Offspring sex ratio as a function of relative patch fecundity per patch in the ‘Global Budding’ (GB, black), ‘Global Random’ (GR, dark grey) and ‘Local Random’ (LR, light grey) selection regimes. Females from the different selection regimes were placed on individual patches (one per patch) with a sterile female from the base population. For each patch, the proportion of offspring produced by the focal female (i.e. from the selection regime) was calculated as the proportion of eggs that hatched and became adult (relative patch fecundity), and her offspring sex-ratio was measured. Each dot represents an individual replicate (i.e., patch from which measurements were taken).

### Sexual conflict

Overall, there was no significant effect of mate number (X^2^_1_=0.024, p = 0.876), male selection regime (X^2^_1_=0.028, p = 0.867), or their interaction (X^2^_1_=0.073, p = 0.788) on the offspring sex-ratio of females from the ancestral population (Figure S3, Table S4). However, the total number of offspring produced was higher when females mated with a male from the ‘Global Budding’, as opposed to the ‘Global Random’, selection regime (X^2^_1_=4.336, p = 0.036; Figure 4, Table S4).

**Figure 4.**
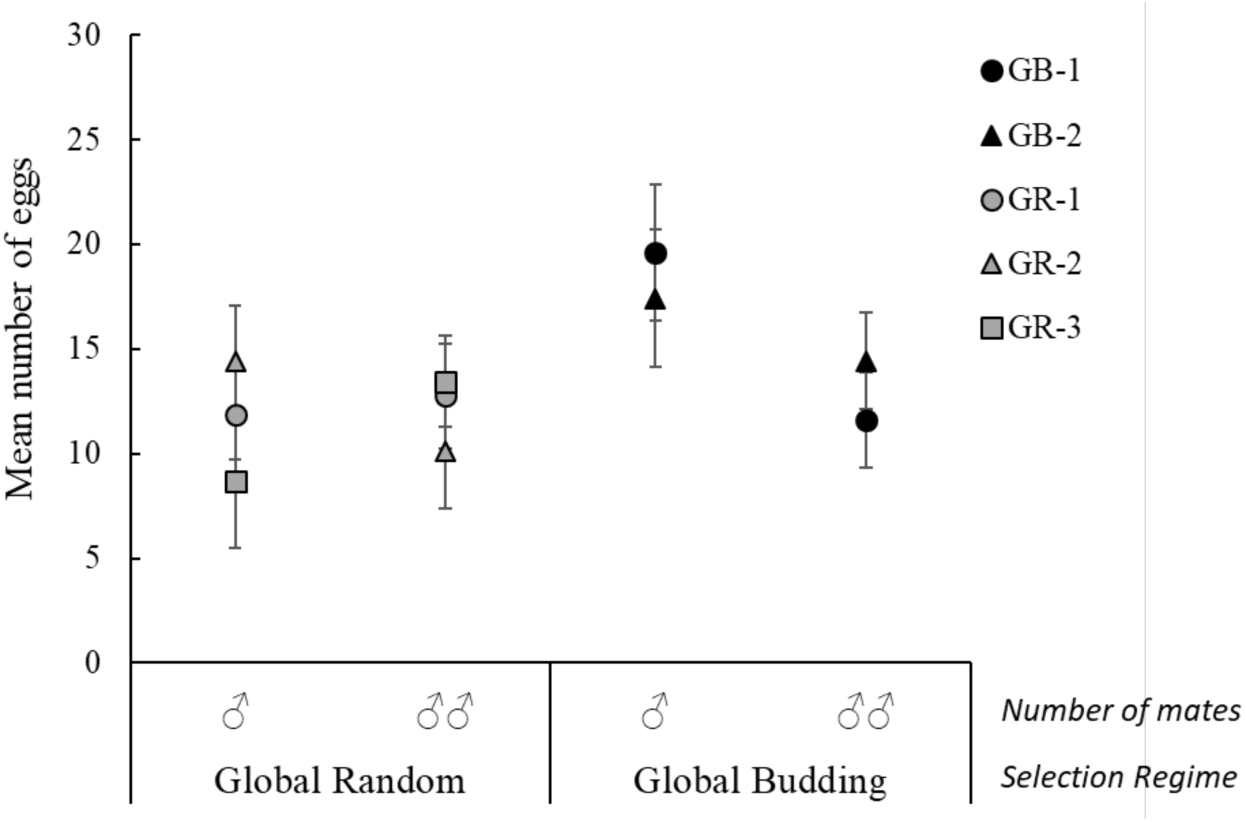
Mean fecundity (± standard error) of females from the ancestral population presented with either one or two males from the ‘Global Budding’ (GB, black) or ‘Global Random’ (GR, dark grey) selection regimes. Means shown for each experimental replicate (different symbols) in each selection regime at generation 33, after two generations in a common environment.

## Discussion

Both sex allocation and sexual conflict responded to selection under different population structures. Sex allocation responses were mainly driven by dispersal type (budding *vs* random), which influences whether interactions occur among kin or unrelated individuals, and not by the scale of competition. Females from the ‘Global Budding’ regime consistently produced more male-biased offspring sex ratios than those from the ‘Global Random’ selection regime. We also found that higher relative patch fecundity was associated with more female-biased offspring sex-ratios across all selection regimes. Finally, when comparing the intensity of sexual conflict, females from the ancestral population mated to males from the ‘Global Budding’ regime had higher fecundity than those mated to males from the ‘Global Random’ regime.

### Sex Allocation

Females from the ‘Global Random’ selection regime produced offspring sex ratios consistent with theory (Taylor and Bulmer 1980). This corroborates previous findings in mites (Macke et al. 2011) and is consistent with sex ratio observations in other haplodiploid and diploid systems (e.g. Herre 1985; Reece et al. 2004; Reece et al. 2008). However, the ‘Local Random’ and the ‘Global Budding’ regimes present offspring sex ratios that differ from theoretical predictions, being more and less female-biased than predicted, respectively (see Table S1). The fact that offspring sex ratios in the ‘Global Budding’ treatment were not as predicted, and that sex ratios in the ‘Local Random’ and ‘Global Random’ regimes were equivalent, suggest that other factors, besides the dispersal type and scale of competition, may be at play. Below, we highlight potential factors that may account for the observed patterns.

First, inbreeding is probably confounded with dispersal type (with high inbreeding expected for the budding dispersal regime) and may be a factor impacting our results. Inbreeding can in itself select for more female-biased offspring sex ratios (Frank 1985; Herre 1985; Chung et al. 2019). If coupled with high levels of juvenile mortality this could, in some cases, result in no males on a patch (West et al. 2002b; Chung et al. 2019), potentially explaining why we lost all 3 replicates of the ‘Local Budding’ and 1 replicate of the ‘Global Budding’ regimes. At the same time, the accrued costs of inbreeding may negate any benefit of female-biased sex ratios in the ‘Global Budding’ regime in the replicates that survived (Greeff 1996). This might be because, in haplodiploids like spider mites, inbreeding costs are expressed mainly in female traits (Tien et al. 2015). As such, there might be selection to augment the production of sons in environments, or patches, with low fecundity and/or high offspring mortality brought about by inbreeding, to ensure female fertilisation (West et al. 2002b; Chung et al. 2019). Thus, it is hard to predict the exact impact of inbreeding in our selection regimes, as we do not have accurate measures of offspring mortality during experimental evolution. However, we do find that females in the ‘Global Budding’ regime produced fewer offspring that became adult, which might be due to higher offspring mortality or lower fecundity (Figure S4, Table S5). These females also produced slightly more male offspring than those from the ‘Global Random’ regime (Figure S5a, Table S5). Finally, it might be that inbreeding reduced variation in the ‘Global Budding’ regime, this way preventing it from being shaped by selection for more female-biased sex ratios. In line with this, the sex ratio in this regime was similar to that observed in natural spider mite populations, which tend to be ∼0.30 (Helle and Sabelis 1985).

Another factor that can further affect sex allocation is variable clutch sizes in a patch. When females with asymmetric fecundities oviposit simultaneously on the same patch, the sons of a more fecund female are subject to more intense competition for mates, as they mostly compete with brothers to mate with sisters; whereas the sons of a less fecund female mostly compete with unrelated males to access unrelated females (Stubblefield and Seger 1990). More fecund females are thus expected to produce more female-biased sex ratios, while less fecund females should produce less female-biased sex ratios. As a result, the patch sex ratio becomes skewed towards that produced by the more fecund females, i.e. a more female-biased sex ratio (Stubblefield and Seger 1990; West 2009). In addition, theory predicts that this adjustment by females should emerge from a differential investment in daughters, while maintaining a constant production of sons (Yamaguchi 1985; Frank 1987). Here, we found that for all selection regimes, the sex ratio declined as the relative fertility of the focal female increased (the same was observed for total patch fecundity, Figure S2), showing that female fecundity and sex-ratio are not independent traits. Furthermore, although not significantly different from the other regimes, ‘Local Random’ females generated the steepest slope, which may suggest a greater tendency for females from this selection regime to adjust their sex allocation in response to fecundity. Coupled with higher overall fecundity observed in this selection regime (Figure S4, Table S4), this may explain why offspring sex ratios are more female-biased than expected. Finally, although son production is not constant across selection regimes (Figure S5a, Table S4), its variation is much lower than that for the number of daughters produced (Figure S5b). Again, this seems to be in line with an effect of clutch size on sex allocation.

Overall, it may be that selection on productivity is driving our results, overriding selection for optimal sex allocation. In the ‘Global Budding’ regime an initial selective sweep for the most productive females may have reduced genetic diversity, which in turn caused higher levels of inbreeding. In the ‘Local Random’ regime (or both ‘random dispersal’ regimes), competition may be selecting for more productive females, which is linked with more female-biased offspring sex ratios.

### Sexual Conflict

It was expected that multiple mating with unrelated males would cause a greater reduction in fecundity, and that they would harass females more, than related males (Pizzari et al. 2015). Accordingly, we found that females mated to males from the ‘Global Random’ regime had lower fecundity than those mated to males from the ‘Global Budding’ regime. This replicates previous findings showing that evolving with kin reduced the level of male inflicted harm (to females) in bulb mites (Lukasiewicz et al. 2017). Other studies have shown that reduced male harm may be a plastic response to the presence of kin (Carazo et al. 2014; Lymbery and Simmons 2017). However, in our experiment, since mating was with unrelated females from the ancestral population, there were no direct cues indicating the presence of kin. In addition, competitor males coming from the same selection regime experienced 2 generations of common garden prior to the experiment which probably reduced relatedness among competitor males. This means that if the response were plastic, then there should be no difference between selection regimes. Thus, reduced harm was most likely an evolved response in our study.

Contrary to expectations (Arnqvist and Rowe 2005), we did not find that multiple mating reduced fecundity in either selection regime. Possibly, the differences in harm inflicted by one or two mates over a single, or two successive five-hour periods respectively, might have been insufficient to detect differences in fecundity between the two treatments. Previous studies done in spider mites found fecundity costs when females were simultaneously exposed to multiple mates for periods of 24h hours (Macke et al. 2012; Rodrigues et al. 2020), or exposed to two mates on multiple occasions during their lifetime (Macke et al. 2012).

Here we only tested the effect of the type of dispersal on sexual conflict. However, the outcome of sexual conflict may also change according to the type of population regulation occurring. Indeed, under local competition, increased competition among relatives is predicted to cancel out the benefits of cooperation (Queller 1992; Taylor 1992; Wilson et al. 1992). This means that sexual conflict might be maintained among related individuals when competition is local (Wild et al. 2011; Pizzari et al. 2015). Yet, despite its general interest, we are not aware of any studies that explicitly test this.

### The interplay between sex allocation and sexual conflict

Evolution under different population structures may simultaneously impact sex allocation and sexual conflict in a non-independent manner (Chapman 2009; Scharer and Janicke 2009). One possibility is that sexual conflict might impact sex allocation if a reduction in female fecundity prevents the production of optimal offspring sex ratios. Our sexual conflict experiment showed that females from the ancestral population mated to males from the ‘Global Random’ regime had the lowest fecundity, suggesting that these males inflict more harm. Yet females from the ‘Global Random’ regime were the most accurate in their sex allocation (Table S1). In addition, ‘Global Random’ females, when mated to ‘Global Random’ males in the sex allocation experiment had higher fecundity (Figure S4, Table S4). This suggests that ‘Global Random’ females may have evolved mechanisms to overcome male harassment or induced harm, as shown in this (Macke et al. 2014) and other (Wigby and Chapman 2004; Michalczyk et al. 2011) systems. Female resistance to harassment may thus be one trait involved simultaneously in the outcome of sexual conflict and sex allocation.

Conversely, sex allocation may also impact sexual conflict through changes in levels of male-male competition; as the sex ratio becomes more male biased so will the intensity of competition. Indeed, evolving with kin has been found to reduce male harm and be associated with more female-biased offspring sex ratios (Lukasiewicz et al. 2017; although the latter was not significantly different from the non-kin evolution treatment). In our sex allocation experiment, sex ratio was the least female-biased in the ‘Global Budding’ selection regime. However, males from this regime inflicted the least harm to females from the ancestral population (sexual conflict experiment), suggesting sex allocation evolution did not result in stronger sexual conflict.

## Conclusions

Our study advances upon previous work investigating the consequences of population structure on sex allocation and sexual conflict. Overall, we show that responses to selection mostly depended on dispersal regime which influences whether interactions are mostly with kin or unrelated individuals. Furthermore, we highlight the need for more studies simultaneously investigating the evolution of sex allocation and sexual conflict, to account for trade-offs between these two traits. To date, we are only aware of one study that considers the evolution of both traits under different population structures (Lukasiewicz et al. 2017), but they only considered relatedness as the variable that differed between treatments. This is despite the fact that non-independence between sex allocation and sexual conflict has been found in a number of systems (Macke et al. 2014; Boulton and Shuker 2015; Boulton et al. 2019) highlighting that associated constraints may cause observed sex ratios or fecundity to differ from predictions.

## Acknowledgements

We would like to thank François Rousset and Elsa Noël for helpful discussion about this manuscript. This work was funded by an ERC consolidating grant (COMPCON GA 725419) to SM, a joint grant from the Agence Nationale de la Recherche and the Fundação para a Ciência e a Tecnologia to Isabelle Olivieri and SM (FCT-ANR/BIA-EVF/0013/2012), a PHC-PESSOA grant (38014YC) to ABD and SM, and a SIRIC Montpellier Cancer Grant (INCa_Inserm_DGOS_12553) funded the irradiator. This is ISEM contribution number 20XX-XXX.

