## Supplementary Materials for "Consequences of population structure for sex allocation and sexual conflict"

**Supplementary Material**

**Table S1. Theoretical predictions for offspring sex ratios under different types of dispersal and scales of competition, and mean observed (± standard error) offspring sex ratios in the different selection regimes.** Here we present predictions for offspring sex-ratios (measured as the proportion of sons) for haplodiploid organisms obtained using several models of sex allocation. Taylor and Bulmer (1980) derive a result for individual dispersal in populations where females mate before dispersal, dispersal is complete, and there is no population regulation prior to dispersal (Taylor and Bulmer 1980). Note, this model (Taylor and Bulmer 1980) advances on Hamilton's original model (Hamilton 1967) providing explicit predictions for haplodiploid species. Thus, it is analogous to the 'Global Random' selection regime (GR, with two foundresses, n=2). Herre (1985) derived a similar model, with the exception of having population regulation before dispersal, which is analogous to the 'Local Random' selection regime (LR, with n=2) (Herre 1985). Gardner et al. (2009) derived a model of budding dispersal with no population regulation which we use to approach the 'Global Budding' selection regime (GB, with two foundresses, n=2; complete budding dispersal dB=1; no migration between patches after budding, m=0) (Gardner et al. 2009). Gardner et al (2009) can also be used to approach the other two regimes. Indeed, full budding dispersal and migration (dB=1, m=1) recovers the result from Taylor and Bulmer (1980), while no budding dispersal but full migration recovers the result from Herre (1985) (thus analogous to the GR and LR selection regimes, respectively). Note that the Gardner et al. (2009) model is expected to break down when considering local competition and no budding dispersal (dispersal dB $\rightarrow$ 0, migration m$\rightarrow$ 0, equivalent to the 'Local Budding' selection regime, LB). Under these conditions there is no genetic mixing between lines for selection to act. Observed offspring sex ratios were defined as the mean offspring sex-ratio of females from the different selection regimes, after 33 generations of selection and two generations in a common environment, in the ‘Sex allocation in response to patch fecundity’ assay.

| | **Budding dispersal** | | **Random dispersal** | |
|---|---|---|---|---|
| | *Predicted* | *Observed* | *Predicted* | *Observed* |
| **Local competition** | na | na | 0.42 | 0.241 ± 0.022 |
| **Global competition** | → 0 | 0.296 ± 0.031 | 0.21 | 0.192 ± 0.020 |

**Table S2. Results of experiment measuring the effect of X-ray irradiation on *T. urticae* survival, fecundity and egg viability.** Groups of thirty *T. urticae* adult females were placed on a bean leaf fragment (16 $cm^2$) on moistened cotton wool in a plastic Petri dish (9 cm diameter). Each group was then irradiated at 0 (control), 10, 25, 50 or 100 Gy, with a dose rate of 2,7 Gy/min using a Xstrahl® XenX pre-clinical irradiator at the Institute of Cancer Research, Montpellier (IRCM). Petri dishes were checked daily for female survival, fecundity (number of eggs laid) and hatchability over the following 6 days. Note that due to logistical constraints, the results in Table S2 correspond to a single replicate (i.e. a single Petri dish containing 30 adult females) per irradiation dose. Results shown refer to measures taken 6 days after irradiation.

| X-ray dose (Gy) | Mortality (%) | Number of eggs produced | Proportion of eggs hatching |
|---|---|---|---|
| **0** | 0 | 457 | 19.5 |
| **10** | 0 | 490 | 7.5 |
| **25** | 0 | 567 | 2.3 |
| **50** | 26.7 | 394 | 2 |
| **100** | 3.3 | 427 | 0 |

**Table S3. Description of the statistical models used for data analysis in each experiment.** Sample size corresponds to the total number of individual replicates (i.e., number of patches from which measurements were taken) included in each analysis. "Maximal model" gives the complete set of explanatory variables (and their interactions) included in the model (note: "*" represents both the interaction between two explanatory variables and their individual effects). "Minimal model" gives the model containing only the variables and the interactions that were statistically significant. Round brackets indicate that the variable was included as a random factor. Square brackets indicate the error structure used ("bb": beta-binomial, "bbI": binomial, accounting for zero inflation; "qp": quasi-poisson, "qpI": quasi-poisson, accounting for zero inflation; "nb": negative binomial). "♂": number of sons; "♀": number of daughters; Generation, "gen": the generations at which the variable of interest was measured across selection regimes during experimental evolution (generations 12, 17, 20 and 31), (note that Generation was analysed as a covariate and was log transformed to improve the fit of the model). Selection Regime, "SelReg": 'Global Budding' (GB), 'Global Random' (GR) and 'Local Random' (LR) (note that in models no. 5 and 6, "SelReg" refers to 'Global Random' (GR) and 'Local Random' (LR) only); Replicate, "rep": experimental replicate; "day": the day when different replicates of the experiment were tested; Number of females, "no. females": the number of females present in a patch (1 or 2) where measurements were taken; Total Patch fecundity, "TPF": the total number of eggs laid by one fertile focal and one sterilised female together on a patch; Relative Patch Fecundity, "RPF": the total number of offspring produced by the focal fertile female divided by the total number of eggs laid by the two females (focal and sterile) present on the same patch; Number of mates, "no. mates": the number of times a female was exposed to a male for 5 hours (single mate or double mates); "box": the container in which several individual replicates were maintained. Model no: 1 = offspring sex ratio during experimental evolution, 2 = offspring sex ratio in a common environment, 3 = offspring sex ratio of the focal female in response to total patch fecundity, 4 = offspring sex ratio of the focal female in response to relative patch fecundity, 5 = offspring sex ratio in the “Sexual conflict” experiment, 6 = total fecundity in the “Sexual conflict” experiment, 7 = total number of adult offspring produced by the focal female in the “Sex allocation in response to patch fecundity” experiment, 8 = total number of sons produced by the focal female in the r”Sex allocation in response to patch fecundity” experiment , 9 = total number of daughters produced by the focal female in the “Sex allocation in response to patch fecundity” experiment. [a] includes all individual replicates measured each generation (generations 12, 17, 20, 31); [b] includes all individual replicates measured after one generation in a common environment (generation 31 + 1), in patches with one or two females and excludes experimental replicate LR-1 due to a lack of individual replicates; [c] includes females that were alive on day 4 and produced offspring and excludes experimental replicates GR-1 and LR-1 due to a lack of individual replicates; [d] only includes individual replicates in which fecundity was higher than zero; [e] only includes individual replicates in which females were alive on day six; [f] includes females that were alive on day 4 and excludes experimental replicates GR-1 and LR-1 due to a lack of individual replicates.

| Model no. | Var. of interest | Response variable | Sample size | Maximal model | Minimal model | R subroutine [err struct.] |
|---|---|---|---|---|---|---|
| 1 | sex-ratio | cbind(♂,♀) | 432 to 384/Gen[a] | log(gen)*SelReg +(rep)+(day) | SelReg +(rep)+(day) | glmmTMB [bb] |
| 2 | sex-ratio | cbind(♂,♀) | 504[b] | no. females*SelReg +(rep) | SelReg +(rep) | glmmTMB [bbI] |
| 3 | sex-ratio | cbind(♂,♀) | 169[c] | TPF*SelReg +(rep) | Eggs+SelReg +(rep) | glmmTMB [bb] |
| 4 | sex-ratio | cbind(♂,♀) | 169[c] | RPF*SelReg +(rep) | RF+SelReg +(rep) | glmmTMB [bb] |
| 5 | sex-ratio | cbind(♂,♀) | 133[d] | no. mates*SelReg +(rep)+(box) | (rep)+(box) | glmmTMB [bb] |
| 6 | total patch fecundity | total number of eggs | 123[e] | no. mates*SelReg +(rep)+(box) | SelReg (rep)+(box) | glmmTMB [qp] |
| 7 | number of offspring | total number of adult offspring | 176[f] | SelReg +(rep) | SelReg +(rep) | glmmTMB [qpI] |
| 8 | number of sons | total number of sons | 169[c] | SelReg +(rep) | SelReg +(rep) | glmmTMB [nb] |
| 9 | number of daughters | total number of daughters | 169[c] | SelReg +(rep) | SelReg +(rep) | glmmTMB [qpI] |

**Table S4. Results obtained from each statistical analysis. "**Df" indicates the degrees of freedom: "$\chi^2$" provides the Chi-square value obtained in each analysis: "Selection regime": 'Global Budding' (GB), 'Global Random' (GR) and 'Local Random' (LR) (note that in models no. 5 and 6, "SelReg" refers to 'Global Random' (GR) and 'Local Random' (LR) only); "Generation": the generations at which the variable of interest was measured across selection regimes during experimental evolution (12, 17, 20 and 31); "Number of females": the number of females present in a patch (1 or 2) where measurements were taken; "Total Patch fecundity": the total number of eggs laid by the focal and sterilised females together on a patch; "Relative Patch Fecundity": the number of offspring produced by the focal female divided by the total number of eggs laid by the two females (focal and sterile) present on the patch; "Number of mates": the number of males a female was exposed to for 5 hours (one or two mates). Model no: 1 = offspring sex ratio during experimental evolution, 2 = offspring sex ratio in a common environment, 3 = offspring sex ratio of the focal female in response to total patch fecundity, 4 = offspring sex ratio of the focal female in response to relative patch fecundity, 5 = offspring sex ratio in the "Sexual conflict" experiment, 6 = total fecundity in the "Sexual conflict" experiment, , 7 = total number of adult offspring produced by the focal female in the "Sex allocation in response to patch fecundity" experiment, 8 = total number of sons produced by the focal female in the "Sex allocation in response to patch fecundity" experiment , 9 = total number of daughters produced by the focal female in the "Sex allocation in response to patch fecundity" experiment. Statistically significant terms in models are represented in bold.

| Model no. | Var. of interest | Explanatory var. | Df | χ2 | P value | Figure |
|---|---|---|---|---|---|---|
| 1 | sex-ratio | Selection Regime x Generation | 2 | 4.351 | 0.114 | 2a |
| | | **Selection Regime** | **2** | **14.046** | **<0.001** | |
| | | Generation | 1 | 2.229 | 0.135 | |
| 2 | sex-ratio | Selection Regime x Number of females | 2 | 4.114 | 0.128 | 2b |
| | | **Selection Regime** | **2** | **11.845** | **0.003** | |
| | | Number of females | 1 | 0.9449 | 0.331 | |
| 3 | sex-ratio | Selection Regime x Total Patch Fecundity | 2 | 0.555 | 0.757 | S2 |
| | | **Selection Regime** | **2** | **9.015** | **0.011** | |
| | | **Total Patch Fecundity** | **1** | **5.366** | **0.021** | |
| 4 | sex-ratio | Selection Regime x Relative Patch Fecundity | 2 | 2.548 | 0.28 | 3 |
| | | **Selection Regime** | **2** | **10.9** | **0.004** | |
| | | **Relative Patch Fecundity** | **1** | **6.87** | **0.009** | |
| 5 | sex-ratio | Number of mates x Selection Regime | 1 | 0.073 | 0.788 | S3 |
| | | Number of mates | 1 | 0.024 | 0.876 | |
| | | Selection Regime | 1 | 0.028 | 0.867 | |
| 6 | total fecundity | Number of mates x Selection Regime | 1 | 0.408 | 0.523 | 4 |
| | | Number of mates | 1 | 1.62 | 0.203 | |
| | | **Selection Regime** | **1** | **4.336** | **0.036** | |
| 7 | number of offspring | **Selection Regime** | **2** | **18.06** | **<0.001** | S4 |
| 8 | number of sons | **Selection Regime** | **2** | **8.365** | **0.015** | S5 |
| 9 | number of daughters | **Selection Regime** | **2** | **10.196** | **0.006** | S5 |

**Table S5.** ***A posteriori*** **contrasts of significant explanatory variables**. A *posteriori* contrasts with Bonferroni corrections were done to interpret the significant effect of selection regime. "Z" = z-scores; "Selection regime": Global Budding (GB), Global Random (GR) or Local Random (LR). Model no: 1 = offspring sex ratio during experimental evolution, 2 = offspring sex ratio in a common environment, 3 = offspring sex ratio of the focal (fertile) female in response to total patch fecundity, 4 = offspring sex ratio of the focal (fertile) female in response to relative patch fecundity, 7 = total number of adult offspring produced by the focal female in the "Sex allocation in response to patch fecundity" experiment, 8 = total number of sons produced by the focal female in the "Sex allocation in response to patch fecundity" experiment , 9 = total number of daughters produced by the focal female in the "Sex allocation in response to patch fecundity" experiment. Statistically significant contrasts are represented in bold (* marginally significant).

| Model no. | Var. of interest | Comparison | Z | P value | Figure |
|---|---|---|---|---|---|
| | | **GB vs GR** | **-3.741** | **<0.001** | |
| 1 | sex-ratio | GR vs LR | 1.554 | 0.361 | 2a |
| | | GB vs LR | -2.289 | 0.066 | |
| | | **GB vs GR** | **-3.384** | **0.002** | |
| 2 | sex-ratio | GR vs LR | -1.597 | 0.3776 | 2b |
| | | GB vs LR | 1.53 | 0.331 | |
| | | **GB vs GR** | **-2.963** | **0.009** | |
| 3 | sex-ratio | GR vs LR | 1.774 | 0.228 | S2 |
| | | GB vs LR | -1.366 | 0.516 | |
| | | **GB vs GR** | **-3.298** | **0.003** | |
| 4 | sex-ratio | GR vs LR | 1.814 | 0.209 | 3 |
| | | GB vs LR | -1.685 | 0.276 | |
| | | **GB vs GR** | **3.523** | **0.001** | |
| 7 | number of offspring | GR vs LR | 0.513 | 1.000 | S4 |
| | | **GB vs LR** | **4.051** | **< 0.001** | |
| | | **GB vs GR** | **-2.634** | **0.025** | |
| 8 | number of sons | GR vs LR | 2.371 | 0.053* | S5 |
| | | GB vs LR | -0.437 | 1 | |
| | | **GB vs GR** | **2.182** | **0.015** | |
| 9 | number of daughters | GR vs LR | 0.213 | 1.000 | S5 |
| | | **GB vs LR** | **2.975** | **0.009** | |

**Figure S1. Schematic representation of the protocol for exposure to a common environment prior to trait measurements in all selection regimes.** Sex allocation in a common environment: At generation 31, ninety-six mated daughters were haphazardly chosen from the 48 patches within each selection regime and placed on a large leaf patch (rectangles) where they laid eggs together. Fourteen days later the offspring on these patches emerged as adults and mated amongst themselves (Generation 31 + 1). After mating, these females were placed on individual leaf patches (squares) where sex allocation measurements were done (see detailed protocol in the main text). Sex allocation in response to patch fecundity: At generation 33, ninety-six mated daughters were haphazardly chosen from the 48 patches within each selection regime and placed on a large leaf patch where they laid eggs together, developed until adulthood and mated. This process was repeated for a second generation (96 mated female offspring from the first generation were placed together on a large leaf patch to lay eggs and offspring to emerge, develop and mate). At the same time, 3 replicate groups of 96 adult mated females from the ancestral population were placed together on a large bean leaf patch to generate sterile females, also over 2 generations (see details in the main text and Table S2). After these two generations in a common environment (Generation 33 + 2), their offspring were used to seed the experiment: single females were placed on leaf patches with a sterile (irradiated) female from the ancestral population (see detailed protocol in the main text). Note that for the 'Sexual Conflict' assay juvenile females were taken from the same mating pools as the mated females used to measure patch fecundity, (not shown in the schematic). ("G": generation).

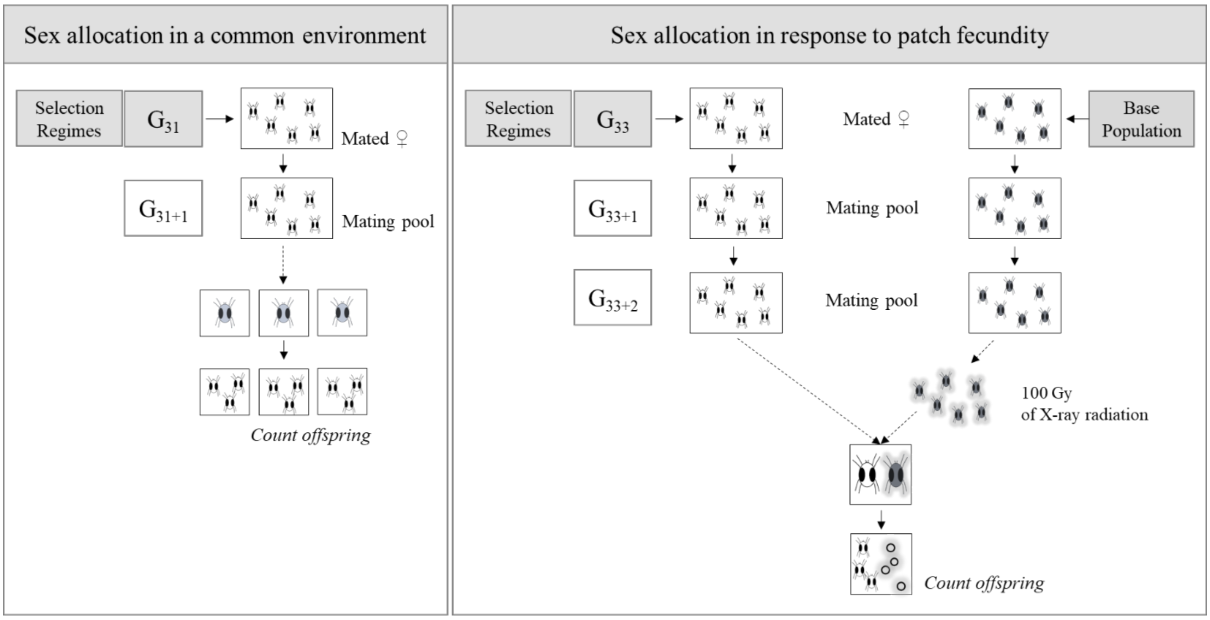
Sex allocation in a common environment
Selection Regimes
G31
Mated ♀
G31+1
Mating pool
Count offspring
Sex allocation in response to patch fecundity
Selection Regimes
G33
Mated ♀
Base Population
G33+1
Mating pool
G33+2
Mating pool
100 Gy
of X-ray radiation
Count offspring

**Figure S2. Offspring sex ratio as a function of total patch fecundity in the 'Global Budding' (black), 'Global Random' (dark grey) and 'Local Random' (light grey) selection regimes.** Females from the different selection regimes were placed on individual patches with a female from the ancestral population that was previously sterilised. On each patch, the total number of eggs laid by both females (total patch fecundity), and the offspring sex-ratio of the focal female (i.e., female from the selection regime) was measured. Each dot represents an individual replicate (the patch from which measurements were taken).

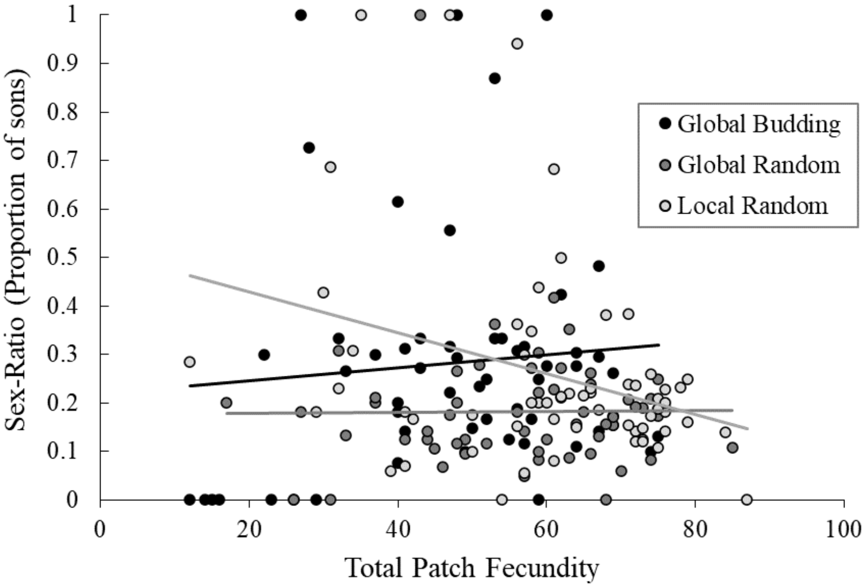

**Figure S3. Mean offspring sex-ratio (± standard error) of females from the ancestral population placed with either one or two mates from the 'Global Budding' (GB, black) or 'Global Random' (GR, grey) selection regimes.** Means are shown for each experimental replicate (different symbols) in each selection regime.

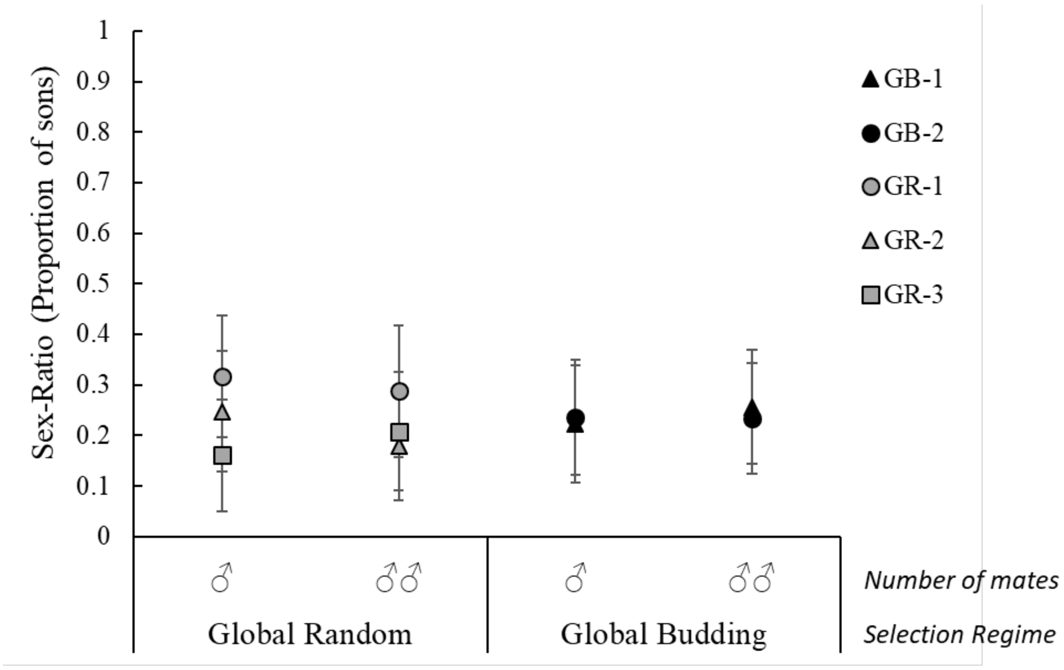

**Figure S4. Total number of adult offspring produced (± standard error) by focal females from the 'Global Budding' (GB, black), 'Global Random' (GR, dark grey) and 'Local Random' (LR, light grey) selection regimes, when sharing a patch with sterilised females from the ancestral population.** Means are shown for each experimental replicate (different symbols) in each selection regime.

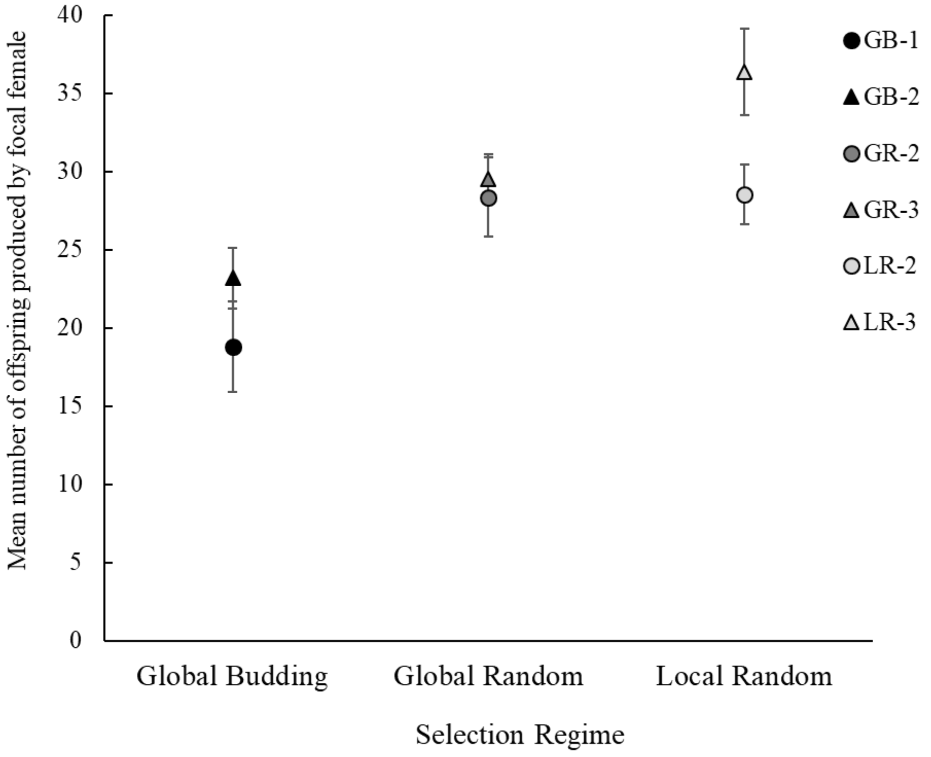

**Figure S5. Total number of adult a) sons and b) daughters (± standard errors) produced by focal females from the 'Global Budding' (GB, black), 'Global Random' (GR, dark grey) and 'Local Random' (LR, light grey) selection regimes, when sharing a patch with sterilised females from the ancestral population.** Means are shown for each experimental replicate (different symbols) in each selection regime.

a) b)

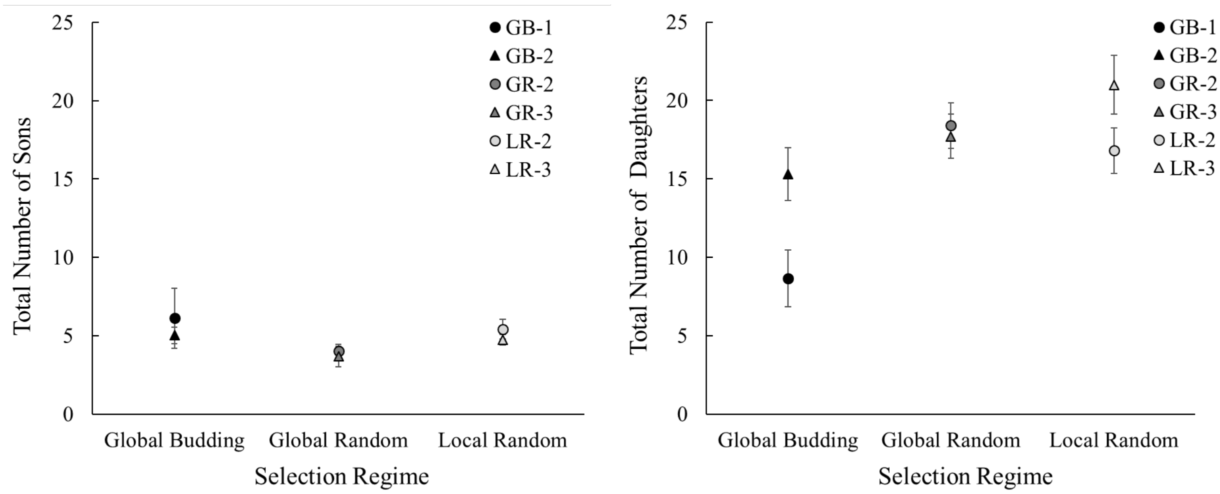
